## Supplementary material for "Effects of blood meal source and seasonality on reproductive traits of *Culex quinquefasciatus* (Diptera: Culicidae)": Figure SM1

**Figure SM1.** Variation of statistical power of the interaction between blood source and seasonality for the fecundity and fertility models across several sample sizes.

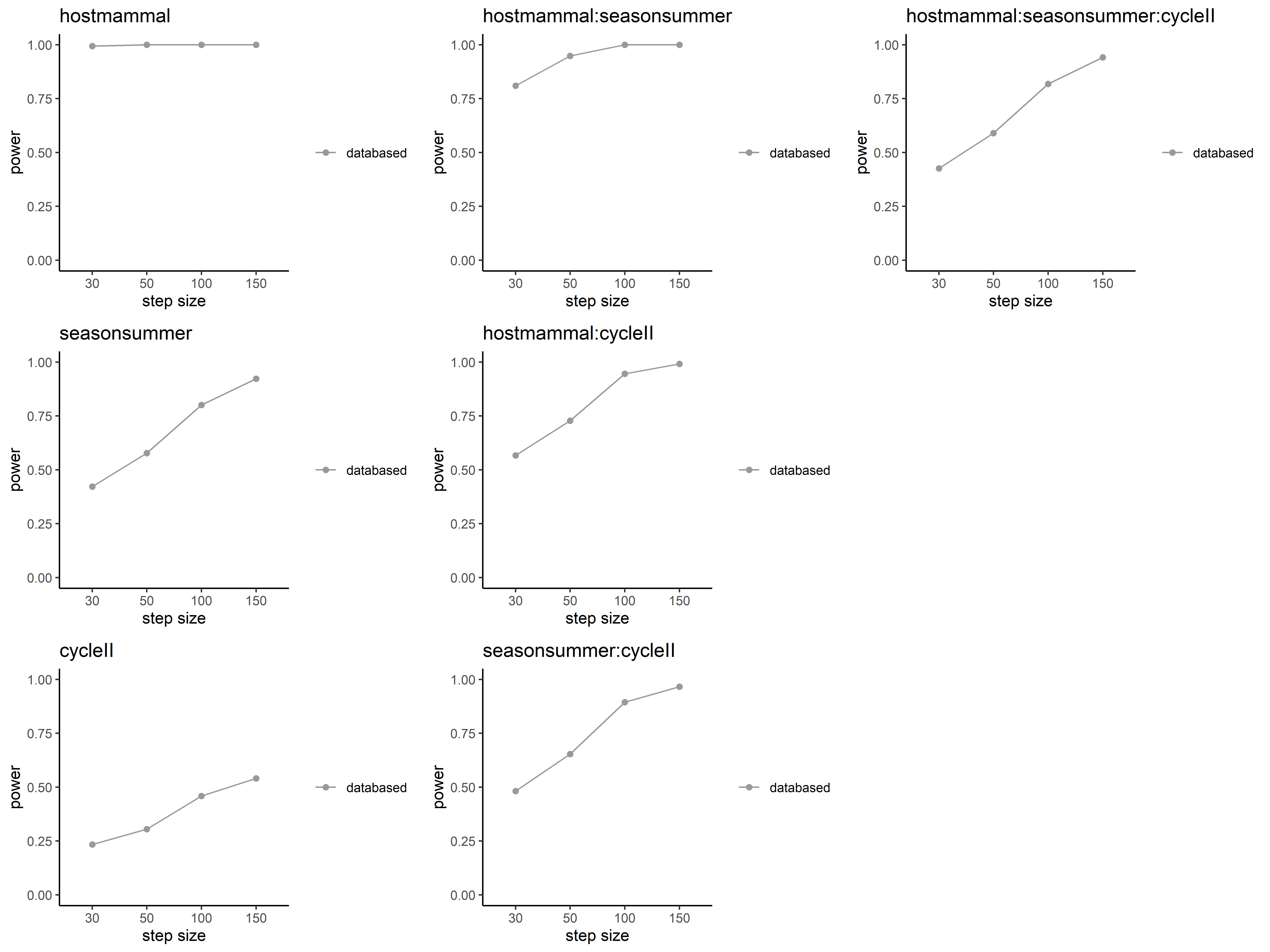

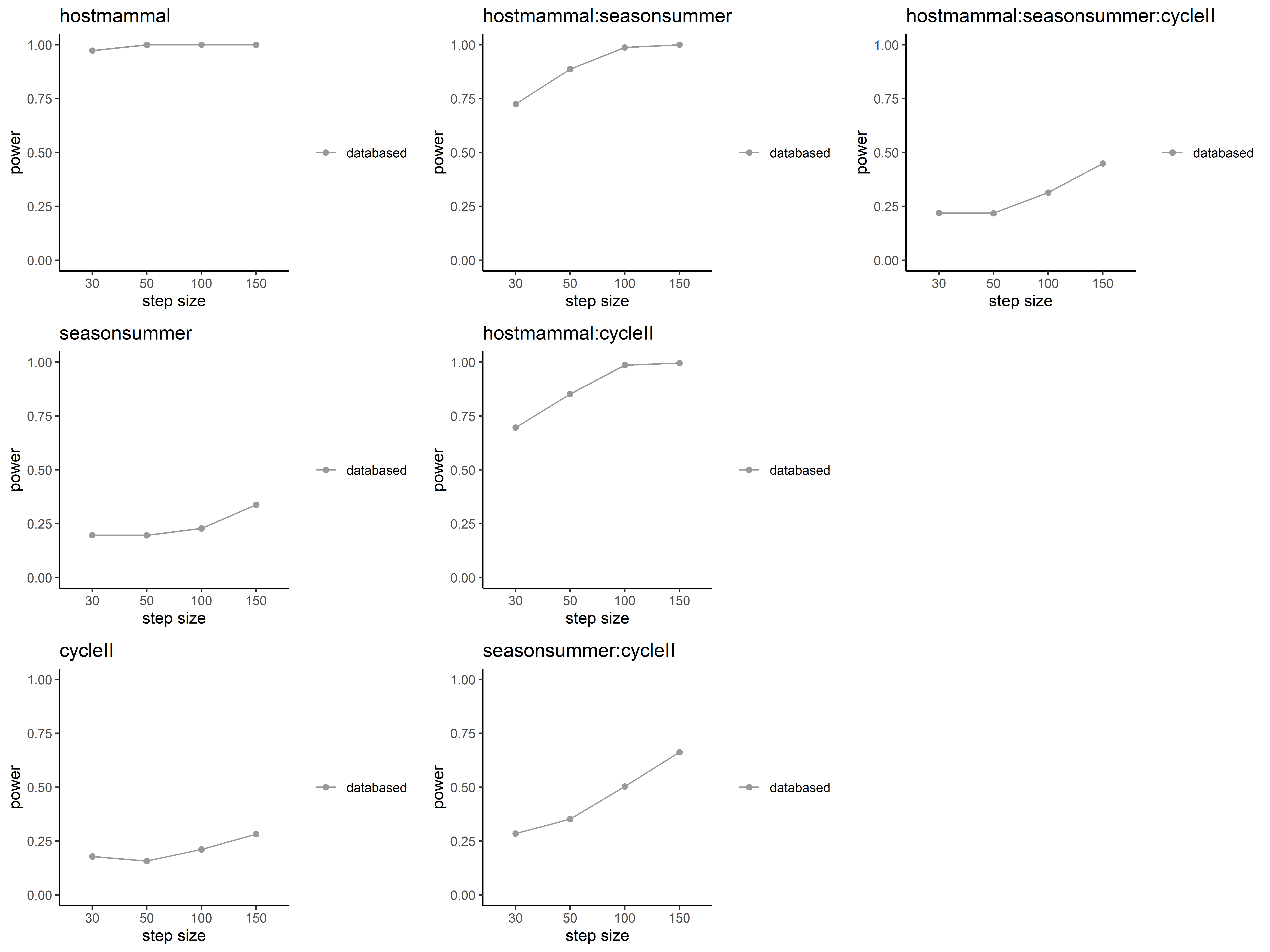

**fecundity (eggs/raft)**

**fertility (larvae/raft)**

**sample size treatment**

**statistical power**
